# On the origin of *R*_2_ orientation dependence angle offsets in white matter

**DOI:** 10.1101/2022.09.16.508261

**Authors:** Yuxi Pang

## Abstract

**Purpose:** to identify the origin of confounding angle offset *ε*_0_ in *R*_2_ orientation dependence and to propose a novel framework for better characterizing anisotropic *R*_2_ in brain white matter (WM).

**Methods:** Anisotropic (*ε*) rather than principal diffusivity direction (Φ) was theorized along axon fiber, with *ε* determined by all eigenvalues and eigenvectors from diffusion tensor. An extra parameter *ε*_0_ was introduced into generalized magic angle effect function to account for any offset in *R*_2_ orientation dependence derived from *T*_2_-weighted image (*b=*0). These dependences referenced by *ε* were compared to those referenced as usual by Φ at *b*-values of 1000 and 2000 (s/mm^2^) on both linear and planar tensor image voxels in WM, based on public domain ultrahigh-resolution (760 *µ*m^3^) Connectome DTI datasets of a healthy young adult brain. A Student’s t-test was used to assess the mean differences and the statistical significance was considered at *P* < 0.05.

**Results:** Fitted *ε*_0_ became zero if referenced by *ε* or nonzero if referenced by Φ, signifying the origin of *ε*_0_. Nonzero *ε*_0_ relied on *b*-values and tensor shapes so did other model parameters, e.g., the amplitude of anisotropic *R*_2_ (1/s) significantly increased when using a higher *b*-value for the linear tensor image voxels, i.e., 3.3±0.1 vs. 1.8±0.1, *P* < 0.01.

**Conclusion:** The origin of *R*_2_ orientation dependence angle offsets has been identified and the combined anisotropic diffusion and transverse relaxation models have fully quantified *R*_2_ orientational anisotropies, thus providing novel insights otherwise unattainable on microstructural alterations in WM.

## 1 INTRODUCTION

A remarkable phenomenon of orientation-dependent MR transverse *R*_2_ (i.e., 1/*T*_2_) relaxation in highly ordered biological tissues, also dubbed magic angle effect (MAE)^1^, has been known for more than half a century.^2^ This specific effect, arising from non-averaging water proton residual dipolar coupling (RDC) in heterogenous local microenvironments,^3, 4^ depends on the direction to an external magnetic field *B*_0_. To date, most basic research and clinical applications on the topic with MR imaging have focused on collagen-rich tissues such as tendon and cartilage.^4-7^ As for the human nervous system, however, MAE remains elusive in the brain white matter (WM)^8-10^ even though it has been reported in the peripheral nervous system.^11, 12^

About 20 years ago, substantial 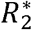 (i.e., 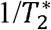) contrast was discovered varying across WM areas at an ultra-high (7T) *B*_0_^13^ and depending largely on fiber tract orientations relative to *B*_0_.^14^ Since then, much effort has been devoted to understanding the relevant mechanisms underlying the measured 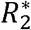 and *R*_2_ orientation dependences at 3T^8, 9, 15-18^ and beyond (≥7T)^10, 19-21^. 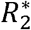 and *R*_2_ contrasts are known originating respectively from gradient-echo and spin-echo based signals. At ultra-high *B*_0_ fields, it was technically challenging to accurately characterize *R*_2_ relaxation because of potential RF heating hazard and/or *B*_1_ inhomogeneity-related incomplete RF refocusing.^10, 22^

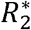 is composed of two components, i.e., 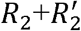, with the former linked to an intrinsic effect arising from local microstructures and the latter to non-local effects from large length-scale field inhomogeneity and macroscopic variation in magnetic susceptibility.^19, 23^ 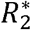 orientation dependence, as elaborated in literature,^19^ comes predominantly from *R*_2_ rather than 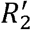. This assessment has been supported by the facts in that not only 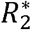 but also *R*_2_ is sorientationally dependent in WM.^9, 15, 24, 25^ For convenience, no distinction will be enforced between 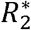 and *R*_2_ when referring to orientation dependence, and the symbol *R*_2_ will be used throughout unless stated otherwise.

In literature, several model functions have been proposed, which were based on microstructural and susceptibility anisotropies in WM, for quantifying orientation-dependent as well as orientation-independent *R*_2_ components, e.g., (1) *a*_0_ + *a*_1_ cos2*ε* + *a*_2_ cos4*ε*,^10^ (2) *b*_0_ + *b*_1_ cos^2^ *ε* + *b*_2_ cos^4^ *ε*,^24^ and (3) *c*_0_ + *c*_1_ sin^2^ *ε* + *c*_2_sin^4^ *ε*.^19, 21, 26^ Unfortunately, these models failed to account for an angle offset *ε*_0_ unexpectedly appeared in *R*_2_ orientation dependences in both adult and neonate WM. More specifically, a small dip between 0° and 30° in anisotropic *R*_2_ profiles in adult WM ^9, 18, 21, 24, 25, 27^ cannot be explained by these prior models and this dip further shifted in neonate WM.^28^ As pointed out lately,^24, 25, 29^ these proposed model functions are mathematically equivalent albeit with different coefficients *a*_*i*_, *b*_*i*_, and *c*_*i*_ (*i* = 0,1,2), thus rendering inconsistent biophysical interpretations on any outcome from these models. It is worth emphasizing that the susceptibility-based relaxation mechanisms^19, 30^ depend quadratically on *B*_0_ while MAE is independent of *B*_0_.^31,32^

Unlike ex vivo studies where WM specimens could be physically repositioned within an MR bore,^10^ the orientation information about an axon fiber in vivo relies exclusively on diffusion tension imaging (DTI) – the only in vivo noninvasive imaging method available to date.^15, 16, 18^ Specifically, principal diffusivity direction (PDD) is assumed to be collinear with a representative axon fiber direction within an image voxel.^33-35^ To our knowledge, such an assumption has never been contested in the past most likely due to the absence of a gold standard. Nevertheless, a number of susceptibility tensor imaging (STI) studies indicate that the principal susceptibility direction deviates to some extent from PDD in WM.^36, 37^ Recently, the most relevant directions associated with anisotropic 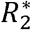 and magnetic susceptibility tensor in WM have been considered collinear with each other,^38^ suggesting that 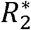 orientation dependence will be shifted when PDD is used as an orientation reference.^19^ Interestingly, it has been further recommended that DTI cannot be regarded as the gold-standard of WM fiber tracts^39^ because PDD does not always reflect exactly actual fiber orientations. This unique perspective is not far away from what has been lately revealed by nanostructure-specific X-ray tomography on axon orientations in WM.^40^

Whether *R*_2_ orientation dependence can be better characterized relies primarily on whether PDD can accurately represent an axon fiber direction. Because of the superior specificity of anisotropic *R*_2_ to highly organized microstructures in WM, it is of high importance to fully quantify the observed *R*_2_ orientation dependences with biologically meaningful and objective model parameters. Thus, the aims of this work were to identify the origin of angle offsets manifested in *R*_2_ orientation dependences, and to propose a novel theoretical framework for better characterizing *R*_2_ orientational anisotropies to attain further insights into microstructural alterations in WM.

## 2 THEORY

### 2.1 An anisotropic diffusion direction

In the human brain WM, thermally driven water molecule translational diffusion is better quantified by a rank-2 tensor ***D***, expressed by a 3×3 symmetric, positive definite matrix.^33^ Because of the symmetry *D*_*ij*_ = *D*_*ij*_ with *i,j* = *x,y,z*, ***D*** has only six independent matrix elements. Based on the standard DTI imaging scheme as stated by Eq. 1,^35^ it requires at least six measurements with a diffusion sensitizing gradient applied successively along noncollinear directions, plus an extra measurement in the absence of the diffusion gradient (i.e., *b*=0) for *SO* normalization, to determine an effective ***D*** from an image voxel.

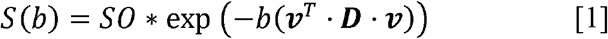

Here, the amount of diffusion weighting is encapsulated by the so-called *b*-value, the diffusion direction is denoted by a unit vector ***v*** in laboratory reference frame (LRF) as shown in Figure 1 with unprimed XYZ axes, and bold face letters symbolize vectors and matrices throughout.

**FIGURE 1.**
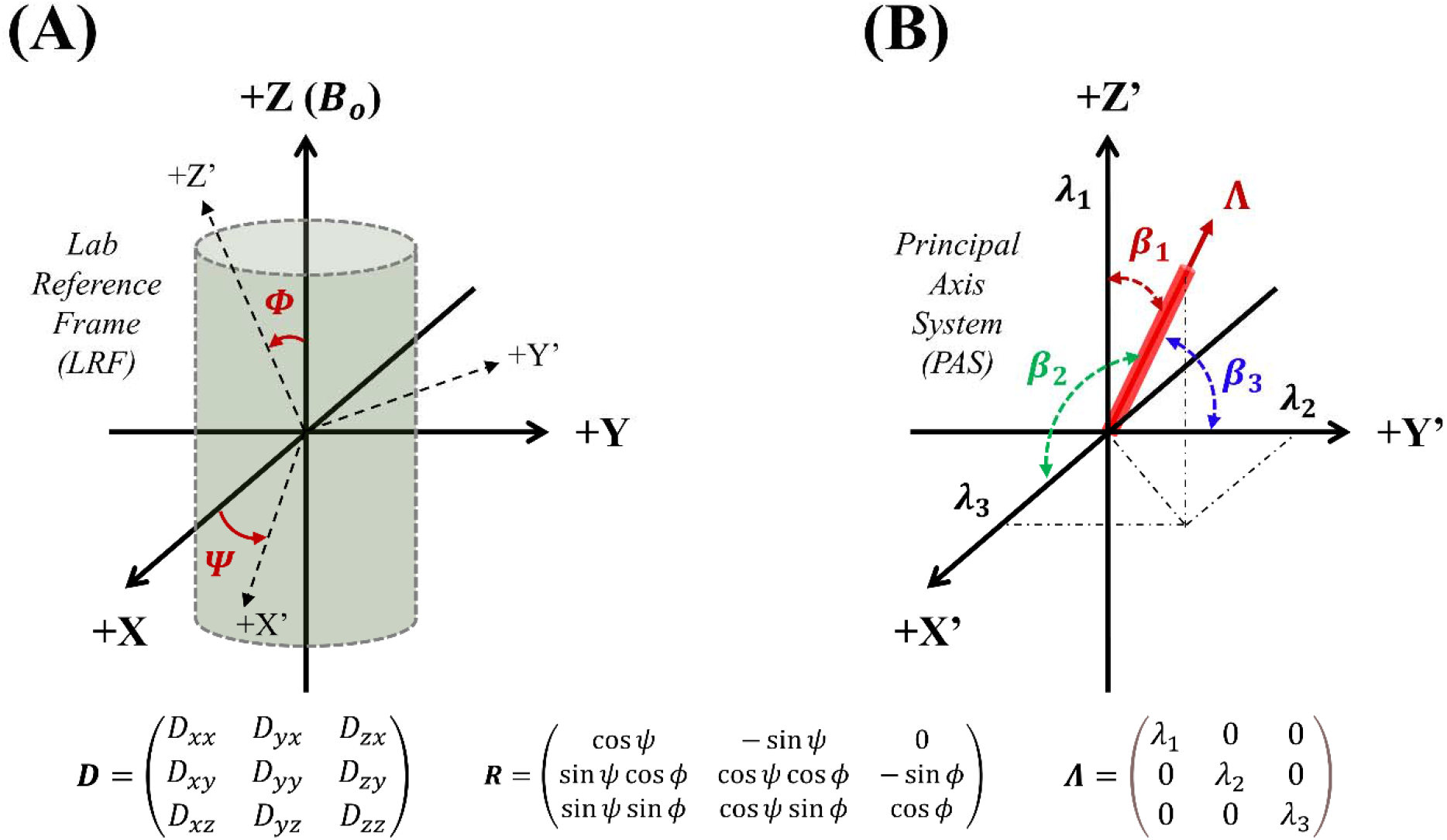
Schematics for transforming diffusion tensor ***D*** from laboratory reference frame (LRF) with unprimed XYZ axis to principal axis system (PAS) with primed XYZ axis. Diagonalized matrix, **Λ** = ***R***^*T*^***DR***, could be further condensed into a vector pointing to anisotropic diffusivity direction as defined by direction angles *βi* (*i = 1,2,3*). ***R*** is rotation matrix and *T* represents transpose operator.

The determined **D** not only reflects microstructural and physiological features underlying the highly organized tissues, but it is also susceptible to physical positions (e.g., rotation) of subjects within an MR scanner.^33^ To derive biologically meaningful and objective diffusion metrics, the determined ***D*** ought to be rotationally transformed into a local reference frame, named principal axis system (PAS) denoted by primed XYZ axes in Figure 1, where ***D*** turns into a diagonalized matrix, i.e., **Λ** = ***R***^*T*^***DR***. The diagonal elements in **Λ** are typically arranged such that *λ*_l_ > *λ*_2_ > *λ*_3_, known as the primary or principal, secondary, and tertiary eigenvalues. Here, ***R*** is a rotation matrix and *T* represents the transpose operator. Note, ***R*** is not known *a priori* but can be determined with Eq. 1 given sufficient diffusion weighting data *S*(*b*).

As schematically shown in Figure 1A, ***R*** may be geometrically constructed by first rotating LRF by an angle Ψ around +Z axis or *B*_0_ direction, and then rotating an additional angle Φ around +X’ axis, leading to an angle Φ between +Z’ in PAS and +Z in LRF.^41^ It is worthy to note that various routes do exist for rotationally transforming from LRF into PAS.^42^ Then, the diagonal elements in ***D*** can be recast in terms of *λ*_*I*_ (*i* = 1,2,3) and associated rotation angles (Φ, Ψ) following an inverse relation of ***D = R*Λ*R***^*T*^; specifically, *D*_*ii*_ (*i* =*x,y,z*) can be expressed with Eq. 2 by assuming an axially symmetric **Λ**.^41^

For a *linear* tensor (i.e., *λ*_l_ > *λ*_2_ ≈ *λ*_3_) as symbolized by a cigar-shaped ellipsoid in Figure 2B, parallel and perpendicular diffusivities are defined respectively as *D*_∥_ = *λ*_l_ and *D*_⊥_ = (*λ*_2_ + *λ*_3_)/2. In contrast, these definitions are different for a *planar* case (i.e., *λ*_l_ ≈ *λ*_2_ > *λ*_3_) as characterized by a plate-shaped ellipsoid in Figure 2E, i.e., *D*_∥_ = *λ*_3_ and *D*_⊥_ = (*λ*_2_ + *λ*_3_)/2. The trace of ***D*** is clearly equal to that of **Λ** as expressed in Eq. 3. In literature, one third of the trace is referred to as mean diffusivity denoted by ⟨*λ*⟩ or ⟨*D*⟩ herein.^33^

**FIGURE 2.**
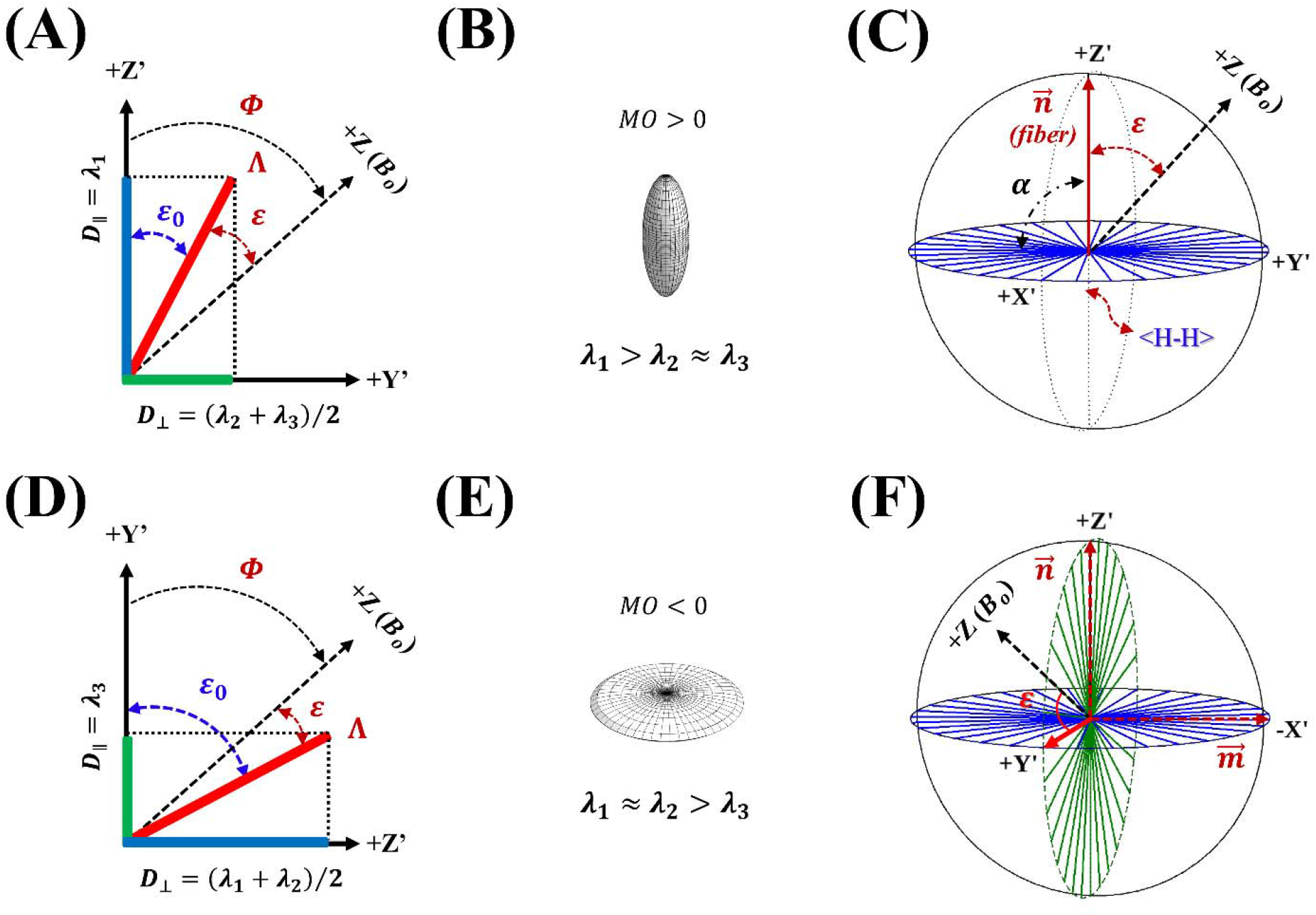
Definitions of orientations relative to *B*_0_ (+Z) in axially symmetric diffusion (2A and 2D) and *R*_2_ relaxation (2C and 2F) models. In diffusion models, *ε*,Φ, and *ε*_0_ denote respectively orientations of anisotropic diffusivity, parallel diffusivity *D* _‖_ defined by primary eigenvalue *λ*_l_ for a linear tensor (2B) or by tertiary eigenvalue *λ*_3_ for a planar tensor (2E), and an angle offset between the first two. Perpendicular diffusivity *D*_⊥_ is the average of two degenerated eigenvalues. For anisotropic *R*_2_ relaxation models, α and *ε* refer respectively to opening angle of the “funnel” model and axon fiber orientation (2C), and these two angles shift by 90° for a fiber crossing case (2F).

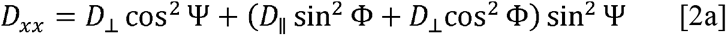

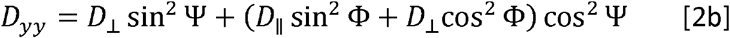

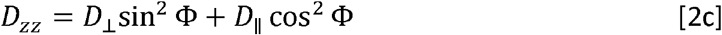

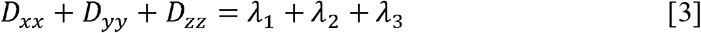

Eigenvectors ***e*_*i*_**, derived from diagonalizing diffusion tensor ***D***, are just as important as the corresponding eigenvalues *λ*_*i*_ (*i* = 1,2,3), which are aligned with PAS orthogonal axes. It is common to assume that principal eigenvector ***e*_1_** points to the direction of an axonal fiber in WM.^34, 35^ However, this common practice appears inconsistent with an advocated basic viewpoint^33^ in that highly ordered microstructures should be exclusively associated with the *anisotropic* component ***D***_*aniso*_ of an effective ***D***.

As shown before, ***D*** can be decomposed into isotropic ***D***_*iso*_ and ***D***_*aniso*_ contributions,^33^ and principal diffusivity *λ*_l_ does not necessarily represent ***D***_*aniso*_ unless it becomes extremely larger than secondary eigenvalue, i.e., *λ*_l_ ≫ *λ*_2_. The magnitude Λ of ***D*** has been defined as the square root of generalized tensor product that is in turn given by three eigenvalues, i.e., 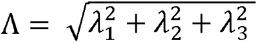. This definition clearly suggests that all diffusion information contained in **Λ**, which has been condensed from ***D***, could be further squeezed into a vector form as demonstrated in Figure 1B, with a unit vector ***e*** = (*e*_*x*_, *e*_*y*_,*e*_*z*_)^*T*^ defined as in Eq. 4.

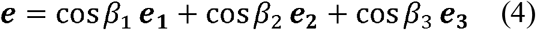

Herein, cos*β*_*i*_ is direction cosine given by *λ*_*i*_/Λ (*i*=1,2,3), and ***e***_*i*_ = (*e*_*ix*_, *e*_*iy*_,*e*_*iz*_)^*T*^an orthonormal column vector in ***R*** that represents direction cosines of each PAS orthonormal axis in LRF.^33^

Since ***D*** comprises both ***D***_*iso*_ and ***D***_*aniso*_ and the former dose not possess any orientation attribute, the direction of the latter ought to be aligned with that of ***D*** or the unit vector ***e*** direction. This unique interpretation agrees well with the prior conclusion in that an effective ***D*** and its component ***D***_*aniso*_ share the same sets of eigenvectors albeit with different eigenvalues offset by mean diffusivity.^33^ Accordingly, an axon fiber orientation should be determined by an anisotropic diffusivity direction (ADD), i.e., *ε* = cos^−l^ *e*_z_, rather than by PDD as usual, i.e., Φ = cos^−l^ *e*_lz_. Thus, it comes as no surprise that previously reported *R*_2_ orientation dependences have manifested an angle offset *ε*_0_ due to an obvious difference between *ε* and Φ. An approximate correction in Φ could be done for an axially symmetric tensor when voxel-based diffusivities or their averages are accessible. In these scenarios, the two direction angles in PAS become complementary, i.e., *β*_l_ + *β*_3_ = 90°. It is also valid for Φ (i.e., cos^−l^ *e*_lz_) and its counterpart (i.e., cos^−l^ *e*_3z_) in LRF. As a result, Eq. 4 could be approximated by Eq. 5a.

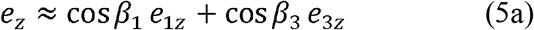

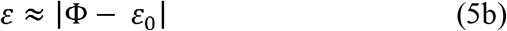

Based on the cosine addition formula in Eq. 5a, *ε* could be estimated with Eq. 5b, where *ε*_0_ has replaced *β*_l_. Accordingly, *ε*_0_ could be determined by tan *ε*_0_ = *D*_⊥_/*D*_∥_ for a linear tensor and by tan *ε*_0_ = *D*_∥_ /*D*_⊥_ for a planar tensor, as schematically illustrated in Figures 2A and 2D. Herein, *D*_∥_ is always associated with the unique diffusivity from an axially symmetric tensor, equal to *λ*_l_ and *λ*_3_ for linear and planar tensors, respectively. Consequently, *R*_2_ orientation dependence referenced by tertiary diffusivity direction (TDD) will be out of phase by 90° when compared to that guided by PDD.

### 2.2 An anisotropic *R*_2_ relaxation model

To expose the *anisotropic R*_2_ relaxation effect hidden in Eq. 1, an image voxel signal *SO* with *b*=0 may be expressed by Eq. 6a,

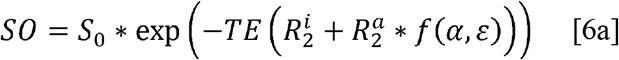

where *TE* denotes a constant spin-echo time in DTI, *S*_0_ is the signal with *TE* = 0, and *R*_2_ has been decomposed, as an effective ***D***, into isotropic 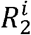 and anisotropic 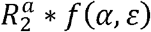 components.^43^ Here, 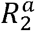 and *f* (*α, ε*) are respectively amplitude and orientation dependence of the anisotropic part. If sufficient orientation information, i.e., *f* (*α, ε*), becomes accessible, it could be more efficient to disentangle the two relaxation components using *T*_2_-weighted images^31, 44^ rather than using conventional time-consuming *T*_2_ maps. Therefore, *SO* in a logarithmic scale, divided by *TE*, could be recast as expressed in Eq. 6b,^45^ where the parenthesized composite term could be considered as an unknown constant, i.e., 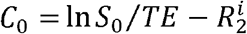.

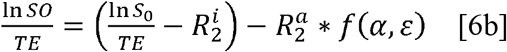

This approximation has been corroborated by the prior observations in that both *S*_0_ and 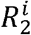, relative to anisotropic *R*_2_, remain essentially unchanged across WM in the human brain.^18^

To parameterize *f* (*α, ε*) for a concentric distribution of bound water RDC around an axon fiber in WM,^46^ an axially symmetric or “funnel” model may be constructed as schematically shown in Figure 2C.^7, 43^ Here, water intramolecular RDC vectors are signified by <H-H>, the angle *ε* is between an axon fiber direction 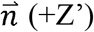 and the main magnetic field *B*_0_ (+Z), and the angle *α* is an opening angle of the funnel. A complex analytical function for this model, as written in Eq. 7a, was given by Berendsen years ago,^2^ with three interdependent coefficients *A*_0_ = 81cos4*α* + 156cos2*α* + 467, *A*_l_ = 180cos4*α* + 432cos2*α* + 156, and *A*_2_ = 315cos4*α* + 180cos2*α* + 81. Recently, this complex function has been reformulated into a compact form as expressed in Eq. 7b, which really highlights the underlying symmetry embodied in the model.^7^

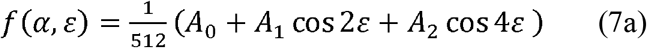

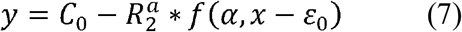

In literature,^18, 24, 25, 28, 29^ Eq. 7a and its variants, as written by a combination of cos^2^ *ε* (or sin^2^ *ε*) and cos^4^ *ε* (or sin^4^ *ε*) terms, have been used for characterizing both isotropic and anisotropic *R*_2_ contributions. In this work, however, *f* (*α, ε*) has been associated entirely with the anisotropic component.

When two axon fibers cross within an image voxel, i.e., 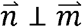, as depicted in Figure 2F, an apparent diffusion tensor could be described by a planar tensor as shown in Figure 2E.^47^ In this work, TDD rather than PDD is used as an unambiguous orientation reference. As a result, *f* (*α, ε*) based on TDD will manifest a 90° shift for either *α* or *ε* compared to that guided by PDD. This is because TDD is aligned with +Y’ in Figure 2F while PDD is collinear with +Z’ in Figure 2C.

## 3 METHODS

### 3.1 Public domain ultrahigh resolution DTI datasets

Public domain ultrahigh resolution (i.e., an isotropic voxel size of 760 *µm*^3^) Connectome DTI datasets of one healthy human brain (age=30 years)^48^ were utilized for validating the proposed theoretical framework. Paired with the associated *b*=0 (s/mm^2^) images, 6 preprocessed data subsets with *b*=1000 (s/mm^2^) and 12 with *b*=2000 (s/mm^2^) were analyzed individually using FSL DTIFIT^49^. Modeled were the following metrics, i.e., eigenvalues *λ*_*i*_ and eigenvectors ***e***_*i*_ (*i*=1,2,3), FA (fractional anisotropy), MO (mode of anisotropy),^50^ *D*_*ij*_ (*i,j* = *x,y,z*), and *T*_2_-weighted signal *SO* with *b*=0. MO could change from 1 to -1, signifying the transition from a perfect linear tensor to an ideal planar tensor. Reconstructed images had a matrix size of 292 by 288 by 192 along respectively left-right, anterior-posterior, and superior-inferior directions.

To evaluate variations of DTI metrics across all data subsets (n=18), a rectangular ROI, 11 pixels wide by 9 pixels high in the corpus callosum in the 97^th^ image slice, was chosen as highlighted by a red arrow in Figure 3D. For each of fitted parameters, an average and its standard deviation were computed. For derivative metrics (e.g., direction angles *β*_*i*_), to determine such descriptive statistics was not straightforward; therefore, the Monte Carlo method^51^ was employed as an alternative. More specifically, 1000 simulations were performed, and each used ROI mean for individual metrics, which was further aggravated by a Gaussian noise with a mean of zero and a standard deviation for the metric within the ROI. Finally, the means and the standard deviations of 1000 simulated derivative metrics were calculated and tabulated in Table 1.

**TABLE 1.** ROI-based DTI metrics and derivatives from the corpus callosum at two *b* values. The measures are tabulated as mean ± SD, and the *P*-value is also provided. Abbreviations: FA, fractional anisotropy; MO, mode of anisotropy.

| Names | $b=1000$ (s/mm<sup>2</sup>) | $b=2000$ (s/mm<sup>2</sup>) | $P$ |
| --- | --- | --- | --- |
| **Ln (SO) (a.u.)** | 6.95 ± 0.07 | 6.97 ± 0.08 | 0.67 |
| $\varepsilon$ (°) | 82.19 ± 1.30 | 79.44 ± 3.15 | 0.06 |
| $\phi$ (°) | 86.26 ± 1.32 | 86.71 ± 0.91 | 0.41 |
| $\varepsilon_0$ (°) | 11.12 ± 1.60 | 17.85 ± 1.87 | < 0.01 |
| $\lambda_1$ ($\mu\text{m}^2/\text{ms}$) | 2.42 ± 0.10 | 1.33 ± 0.04 | < 0.01 |
| $\lambda_2$ ($\mu\text{m}^2/\text{ms}$) | 0.69 ± 0.07 | 0.54 ± 0.05 | < 0.01 |
| $\lambda_3$ ($\mu\text{m}^2/\text{ms}$) | 0.23 ± 0.07 | 0.31 ± 0.04 | < 0.01 |
| $\beta_1$ (°) | 18.05 ± 2.00 | 25.68 ± 2.46 | < 0.01 |
| $\beta_2$ (°) | 74.07 ± 1.57 | 68.28 ± 1.89 | < 0.01 |
| $\beta_3$ (°) | 84.75 ± 1.53 | 77.76 ± 1.42 | < 0.01 |
| **FA** | 0.79 ± 0.03 | 0.63 ± 0.04 | < 0.01 |
| **MO** | 0.76 ± 0.03 | 0.72 ± 0.07 | 0.3 |

**FIGURE 3.**
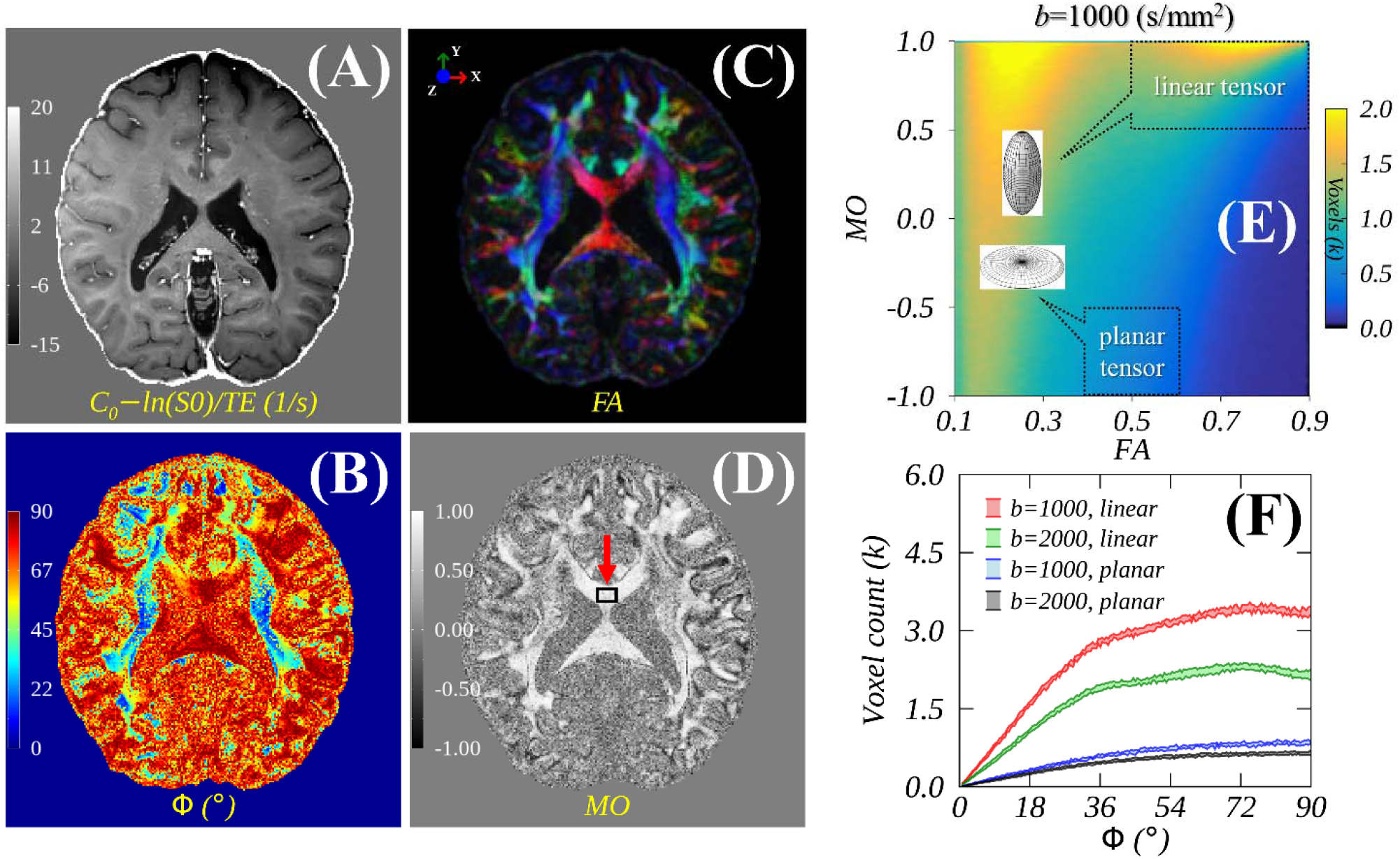
Illustrative parametric maps of *C*_0_ − ln(*SO*)/*TE* or anisotropic *R*_2_ (3A), principal diffusivity orientation Ф (3B), color-coded fractional anisotropy (3C), and mode of anisotropy (3D). Linear and planar tensors were categorized as shown in a 2D (i.e., FA vs. MO) histogram of whole brain image voxels (3E). Orientation-dependent voxel counts are presented from both linear and planar tensor groups at two *b*-values (3F).

### 3.2 Different orientation references

Because of the symmetry in *f* (*α, ε*),^7^ an axon fiber orientation was limited to [0°, 90°] when referenced by the proposed ADD, i.e., *ε* = cos^−l^|*e*_z_| and by the standard PDD for a linear tensor, i.e., Φ = cos^−l^|*e*_lz_| or by TDD for a planar tensor, i.e., Φ = cos^−l^|*e*_3z_|. Image voxels from the whole brain WM associated with linear and planar tensors were categorized by a combination of FA and MO limits as shown in a 2D histogram in Figure 3E. Specifically, the linear tensor group was determined by 0.5 < *FA* < 0.9 and 0.5 < *MO* < 1.0 while the planar tensor group was by 0.4 < *FA* < 0.6 and -1.0 < *MO* < -0.5.

The determined *ε* and Φ were sorted ascendingly from 0° to 90° as usual,^15, 16, 18^ and the corresponding metric was averaged within each predefined data bin to generate an angular dependence profile. The data bin size was 0.5° and the presented orientation-dependent metrics in all plots were sorted according to Φ unless specified otherwise. Sorted by both *ε* and Φ, the scaled *T*_2_-weighted signal intensities, i.e., *y* = *lnSO*/*TE*, were compared. The measured *y* values, i.e., anisotropic *R*_2_ rates, were fitted to Eq. 7, which originated from Eq. 6b except for an extra model parameter *ε*_0_ accounting for an angle offset.^43, 45^

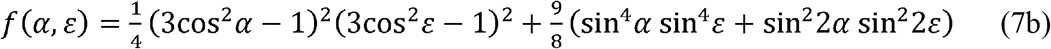

Here, independent parameter *x* was either *ε* or Φ, and *TE* was 75ms used in DTI.^48^ There were four model parameters to be fitted, i.e., 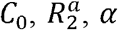, and *ε*_0_, and *y* was dependent parameter. The inputs *x* and y were respectively the averages of the means across all data subsets acquired using the same *b* value. While the input *x* was considered precise, the measurement uncertainty in y was taken from its standard deviation of the means across all data subsets with the same *b* value.

### 3.3 Nonlinear least-squares anisotropic *R*_2_ modeling

A public domain nonlinear least-squares optimization program (http://purl.com/net/mpfit) was used for weighted data modeling with model parameters constrained.^52^ The searching ranges for optimal model parameters were limited as follows: 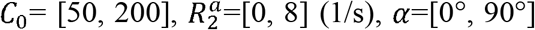, and *ε*_0_=[0°, 45°]. A maximum number of iterations was 1000, divided into five subgroups. To avoid potential local minima during optimization processes, each subgroup started with different initial searching values within the constraints. The goodness of fits was characterized by an adjusted R squared in percent (RSQ%).

Unless otherwise specified, the fitted y was plotted with a red solid line, and primary, secondary, and tertiary diffusivities *λ*_*i*_ and direction angles *β*_*i*_ (*i* = 1,2,3) were respectively represented by red, green, and blue ribbons in all plots. Wherever applied, error bars or half widths of ribbons were determined by the related standard deviations. A Student’s paired t-test with a two-tail distribution was used to assess the mean difference in variants between from *b*=1000 (s/mm^2^) and from *b*=2000 (s/mm^2^), and the statistical significance was considered at *P* <

0.05. Data analysis and visualization were performed using in-house software developed in IDL 8.8 (Harris Geospatial Solutions, Inc., Broomfield, CO, USA).

## 4 RESULTS

### 4.1 Exemplary parametric maps

Figure 3 displays four parametric maps derived from six DTI data subsets (*b* =1000 s/mm^2^) including (3A) *C*_0_ – ln(*SO*)/*TE* or anisotropic *R*_2_ (1/s), (3B) principal diffusivity orientation Φ (°), (3C) color-coded fractional anisotropy, and (3D) mode of anisotropy. A 2D histogram (FA vs. MO) reveals a distribution of all image voxels from whole brain (3E), with two regions highlighted by callout boxes to distinguish the linear and planar tensor groups. Angular dependent voxel counts among different groups are approximately proportional to sinΦ (3F), in agreement with previous findings.^15, 53^ The voxel counts from =1000 (s/mm^2^) are always larger than those from *b* =2000 (s/mm^2^) particularly in the linear tensor group, i.e., red vs. green ribbons.

### 4.2 Exemplary parametric variations

Figure 4 illustrates variations of DTI metrics and derivatives from an ROI as highlighted in Figure 3D, and Table 1 tabulates the summary statistics. Comparing results from *b*=2000 (s/mm^2^) to those from *b*=1000 (s/mm^2^), no significant changes are found for *SO* (*P* = 0.67, 4A) and MO (*P* = 0.3, 4F); in contrast, significant (*P* < 0.01) differences appear in eigenvalues (4C) and direction angles (4D), e.g., 1.33 ± 0.04 vs. 2.42 ± 0.10 for principal diffusivity *λ*_l_ (*µ*m^2^/ms). This finding is also mirrored by a significant decline in FA, i.e., 0.63±0.04 vs. 0.79±0.03 (4E), which is at odds with a previous finding at 1.5T for frontal WM and internal capsule.^54^ The observed discrepancy could be ascribed to an inadequate SNR at a lower *B*_0_ field or a partial volume effect due to a larger voxel size (i.e., 1.56mm x 1.56mm × 10.0mm) in the previous study.

**FIGURE 4.**
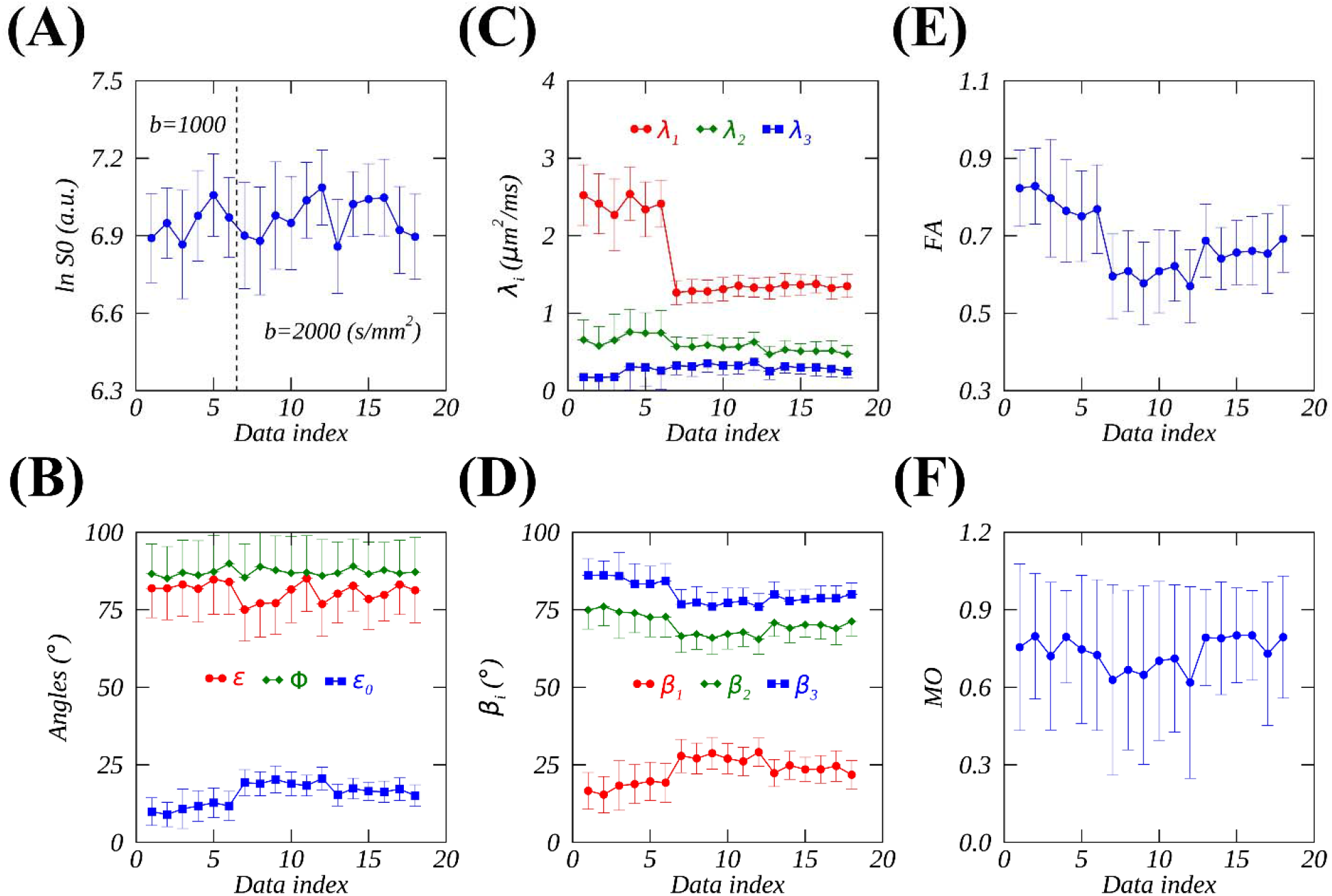
Illustrative DTI metrics and derivatives averaged (mean ± SD) within an ROI in the corpus callosum (see Figure 3D) for all data subsets (n=18). These plots include (4A) *T*_2_-weighted signal *SO* in a logarithmic scale (4A); anisotropic diffusivity orientations (red), principal diffusivity orientations (green), and predicted angle offsets (blue) (4B); primary (red), secondary (green) and tertiary (blue) diffusivities (4C) and direction angles (4D); fractional anisotropy (4E) and mode of anisotropy (4F). Note, the first six data subsets were acquired using *b*=1000 (s/mm^2^) and the rest using *b*=2000 (s/mm^2^), and error bars denote standard deviations (SD).

As shown in Figure 4B, the predicted angle offset (*ε*_0_ in blue) undergoes a significant (*P* < 0.01) increase; however, the changes are insignificant for either principal diffusivity angles (*ϕ* in green, *P* = 0.41) or anisotropic diffusivity angles (*ε* in red, *P* = 0.06). More importantly, *ε* is always smaller than Φ and the gaps become wider at a higher *b* value, implying that an intrinsic different *R*_2_ orientation dependence would appear when using a different *b* value or being guided by a different orientation reference.

### 4.3 *R*_2_ orientation dependences from linear tensor group

Figure 5 presents orientation-dependent DTI metrics derived from data subsets (n=6) with *b*=1000 (s/mm^2^) for those image voxels in the linear tensor group. While diagonal elements *D*_*ii*_ (*i* = *x,y,z*)depend on orientations (5B), their counterparts *λ*_1_ and *β*_*i*_ (*i* = 1,2,3) are rotationally invariant except for *λ*_l_ (5A and 5C). This unexpected slight alteration in *λ*_l_ is also echoed in the predicted *ε*_0_ with an opposite trend due to tan *ε*_0_ = *D*_⊥_/*D*_∥_(5D).

**FIGURE 5.**
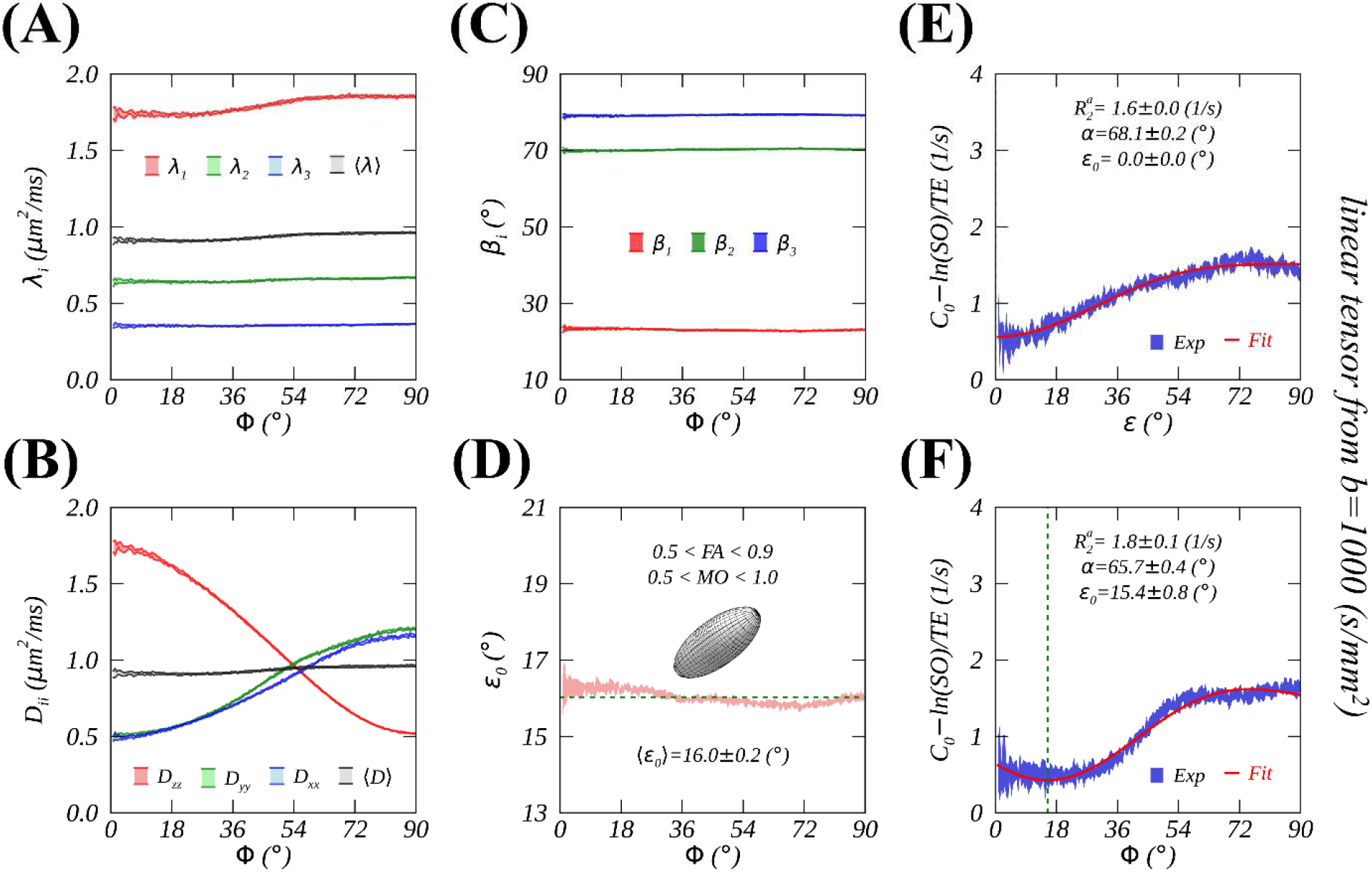
Linear diffusion tensor metrics and *R*_2_ orientation dependences from DTI data subsets (n=6) with *b*=1000 (s/mm^2^). Diffusion metrics include diagonal matrix elements in PAS (5A) and LRF (5B), anisotropic diffusivity direction angles (5C), and predicted angle offsets (5D). *R*_2_ orientation dependence was guided either by anisotropic diffusivity direction *ε* without an angle offset *ε*_0_ (5E) or by principal diffusivity direction Φ with an angle offset *ε*_0_ (5F). Abbreviations: PAS, principal axis system; LRF, laboratory reference frame.

When *R*_2_ orientation dependence was guided by *ε*, there was no angle offset as shown by the fitted *ε*_0_=0 (5E). In contrast, this fitted *ε*_0_ became nonzero (5F) when Φ was used as an orientation yardstick as previously observed.^18, 24, 25^ This angle offset could be essentially removed when a voxel-wise *ε*_0_ correction (i.e., *ϕ* − *ε*_0_) was enforced as demonstrated lately.^43, 45^ These results clearly reveal the origin of previously observed angle offsets in *R*_2_ orientation dependences in WM.

Figure 6 presents the same plots as those in Figure 5 except from data subsets (n=12) with *b*=2000 (s/mm^2^). Overall, the same conclusions could be drawn as those from *b*=1000 (s/mm^2^). Nevertheless, *λ*_1_ became less fluctuated (6A) and eigenvalues came closer indicating a reduced FA, which was further resonated by changes in *D*_*ii*_ (6B), *β*_*i*_ (6C) and predicted *ε*_0_ (6D).

**FIGURE 6.**
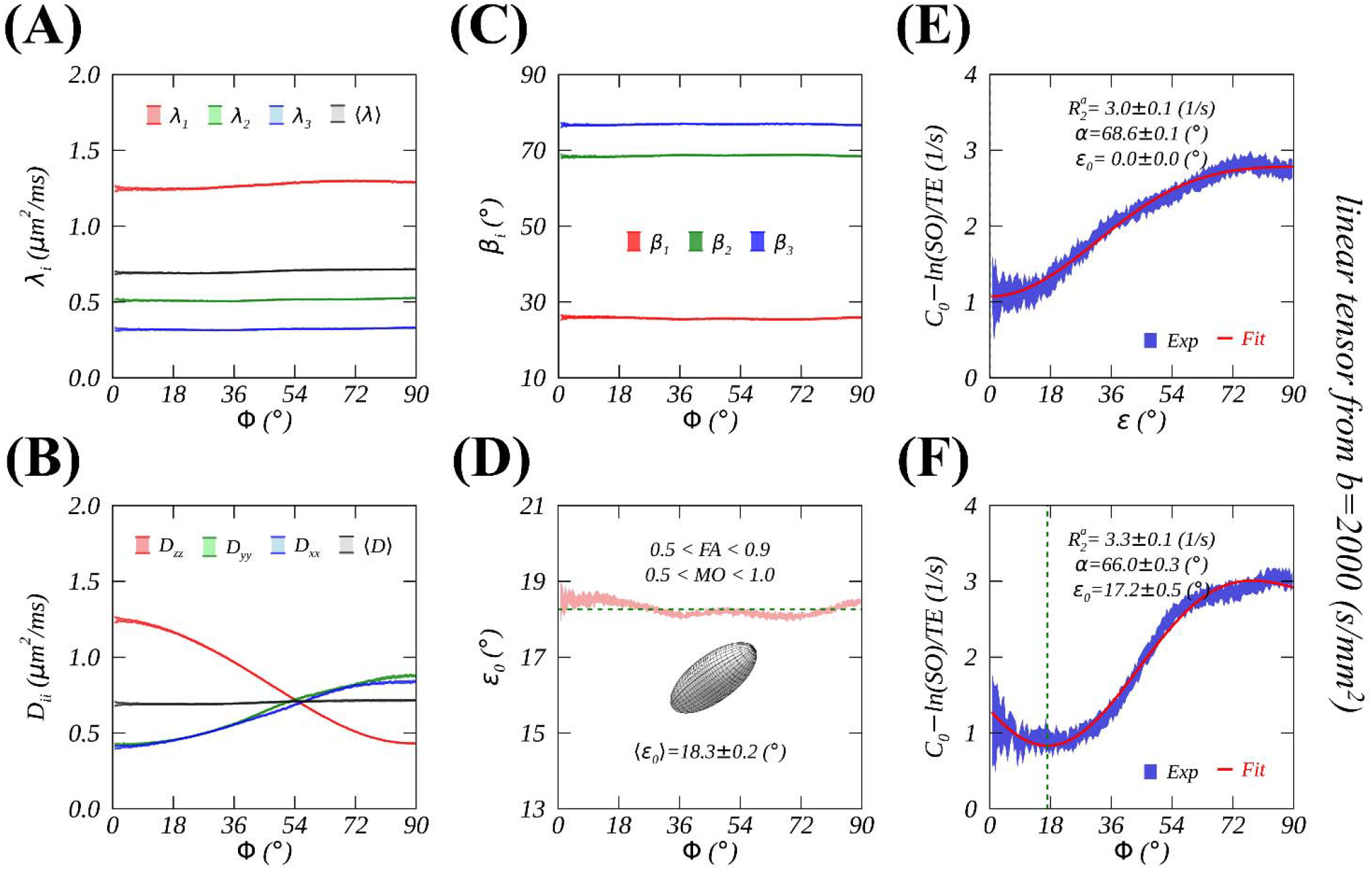
The same plots as Figure 5 except for the results derived from DTI data subsets (n=12) with *b*=2000 (s/mm^2^) containing linear tensor image voxels.

The detailed descriptive statistics are listed in Table 2.

**TABLE 2.** DTI metrics and derivatives in WM at two *b* values. Image voxels from whole brain WM were divided into linear and planar tensor groups. The measures are tabulated as mean ± SD. Angle brackets refer to an average. Abbreviation: RSQ, adjusted R-squared; WM, white matter.

| Names | linear tensor |  | planar tensor |  |
| --- | --- | --- | --- | --- |
| | $b=1000$ (s/mm<sup>2</sup>) | $b=2000$ (s/mm<sup>2</sup>) | $b=1000$ (s/mm<sup>2</sup>) | $b=2000$ (s/mm<sup>2</sup>) |
| $\lambda_1$ ($\mu\text{m}^2/\text{ms}$) | $1.80 \pm 0.05$ | $1.27 \pm 0.02$ | $1.34 \pm 0.01$ | $1.01 \pm 0.00$ |
| $\lambda_2$ ($\mu\text{m}^2/\text{ms}$) | $0.65 \pm 0.01$ | $0.51 \pm 0.01$ | $1.11 \pm 0.01$ | $0.84 \pm 0.00$ |
| $\lambda_3$ ($\mu\text{m}^2/\text{ms}$) | $0.36 \pm 0.00$ | $0.32 \pm 0.00$ | $0.38 \pm 0.00$ | $0.31 \pm 0.00$ |
| $\beta_1$ ($^\circ$) | $23.04 \pm 0.22$ | $25.66 \pm 0.21$ | $77.55 \pm 0.09$ | $76.72 \pm 0.08$ |
| $\beta_2$ ($^\circ$) | $70.21 \pm 0.19$ | $68.58 \pm 0.17$ | $51.60 \pm 0.05$ | $51.60 \pm 0.05$ |
| $\beta_3$ ($^\circ$) | $79.25 \pm 0.14$ | $76.84 \pm 0.12$ | $41.27 \pm 0.07$ | $41.59 \pm 0.06$ |
| $\langle \varepsilon_0 \rangle$ ($^\circ$) | $16.02 \pm 0.17$ | $18.27 \pm 0.16$ | $72.55 \pm 0.14$ | $71.46 \pm 0.12$ |
| $\varepsilon_0$ ($^\circ$) | $15.35 \pm 0.76$ | $17.16 \pm 0.54$ | $0.02 \pm 4.42$ | $2.06 \pm 4.29$ |
| $\alpha$ ($^\circ$) | $65.67 \pm 0.43$ | $65.96 \pm 0.34$ | $26.61 \pm 1.33$ | $25.63 \pm 1.66$ |
| $R_2^a$ (1/s) | $1.77 \pm 0.07$ | $3.30 \pm 0.10$ | $0.68 \pm 0.12$ | $0.86 \pm 0.18$ |
| $C_0$ (a.u.) | $96.73 \pm 0.06$ | $98.13 \pm 0.07$ | $95.49 \pm 0.06$ | $95.49 \pm 0.10$ |
| RSQ (%) | 97.74 | 99.05 | 85.17 | 94.95 |

Although the fitted *α* (°) are consistent across *b* values based on either *ε* or Φ, the fitted amplitude 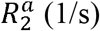 is always significantly (*P* < 0.01) larger when using an increased *b* value. For instance, 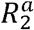 increased from 1.6±0.0 (1/s) at *b*=1000 (s/mm^2^) (5E) to 3.0±0.1 (1/s) at *b*=2000 (s/mm^2^) (6E) when *ε* was used. If Φ was used, the fitted *ε*_0_ slightly increased at a higher *b* value (i.e., 6F vs. 5F) as indicated by an increased average ⟨*ε*_0_⟩ (i.e., 6D vs. 5D). Nevertheless, no angle offset appeared (i.e., 6E vs. 5E), as predicted in Theory section, when the proposed *ε* was used as an orientation reference.

### 4.4 *R*_2_ orientation dependences from planar tensor group

Figures 7 and 8 show the same plots as those in Figures 5 and 6 but from image voxels in the planar tensor group. As seen in the linear case, a higher *b* value led to a decreased FA, indicated by reduced differences among *λ*_*i*_ (i.e., 8A vs. 7A) and *D*_*ii*_ (i.e., 8B vs. 7B). Unlike an increase for its counterpart (i.e., 18.27 ± 0.16° vs. 16.02 ± 0.17°, 6D vs. 5D), the predicted ⟨*ε*_0_⟩ for the planar case became much larger and slightly reduced (i.e., 71.46 ± 0.12° vs. 72.55 ± 0.14°, 8D vs. 7D) with a higher *b* value. Obviously, the predicted ⟨*ε*_0_⟩ failed to correct slightly shifted *R*_2_ orientation dependence profiles guided by Φ (i.e., 7F and 8F). Nonetheless, when referenced by *ε*, these profiles did not show any angle offsets in Figures 7E and 8E, further demonstrating the origin of the angular offsets in image voxels containing crossing fibers.

**FIGURE 7.**
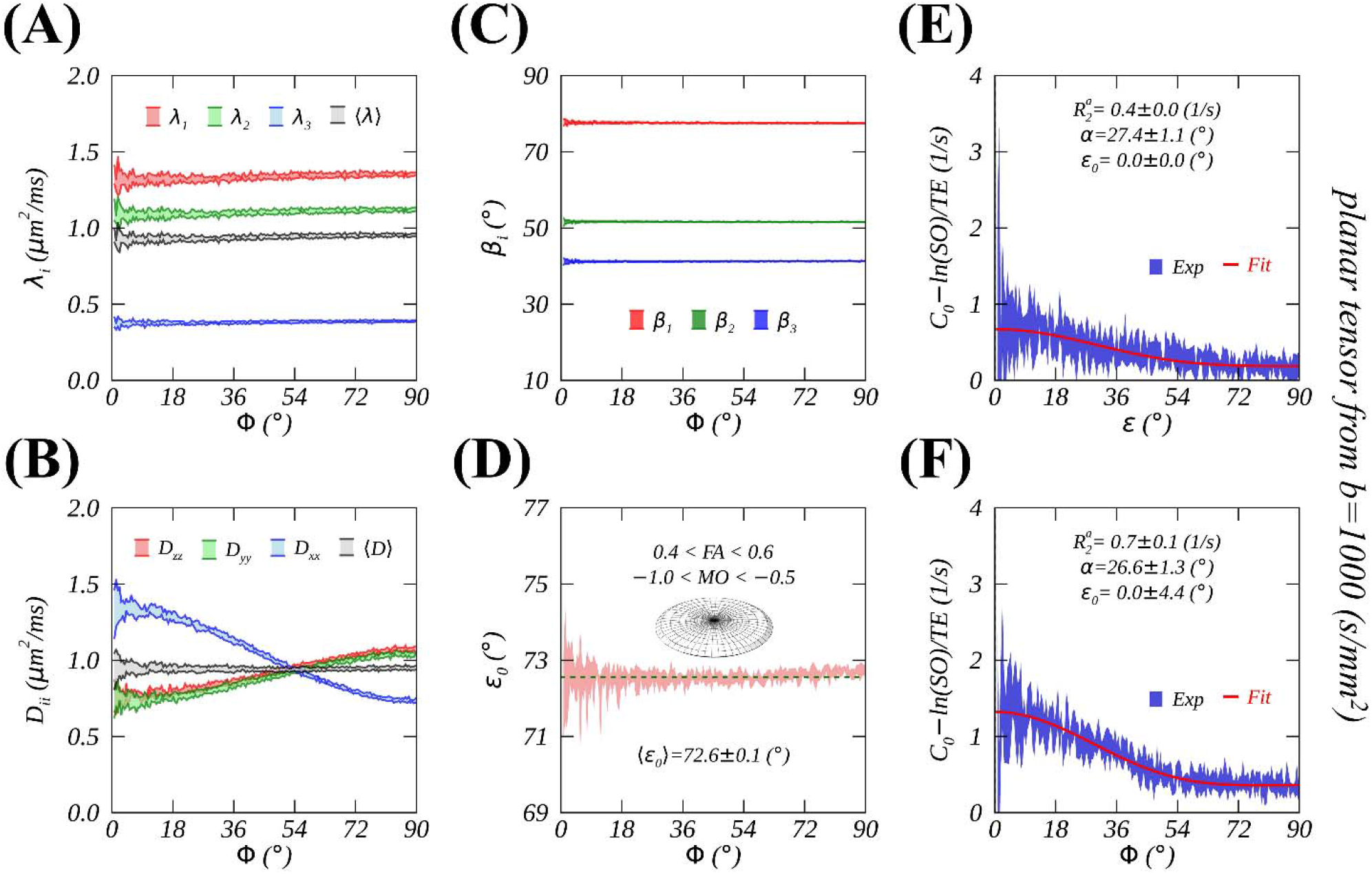
The same plots as Figure 5 except for the results derived from DTI data subsets (n=6) with *b*=1000 (s/mm^2^) containing planar tensor image voxels.

**FIGURE 8.**
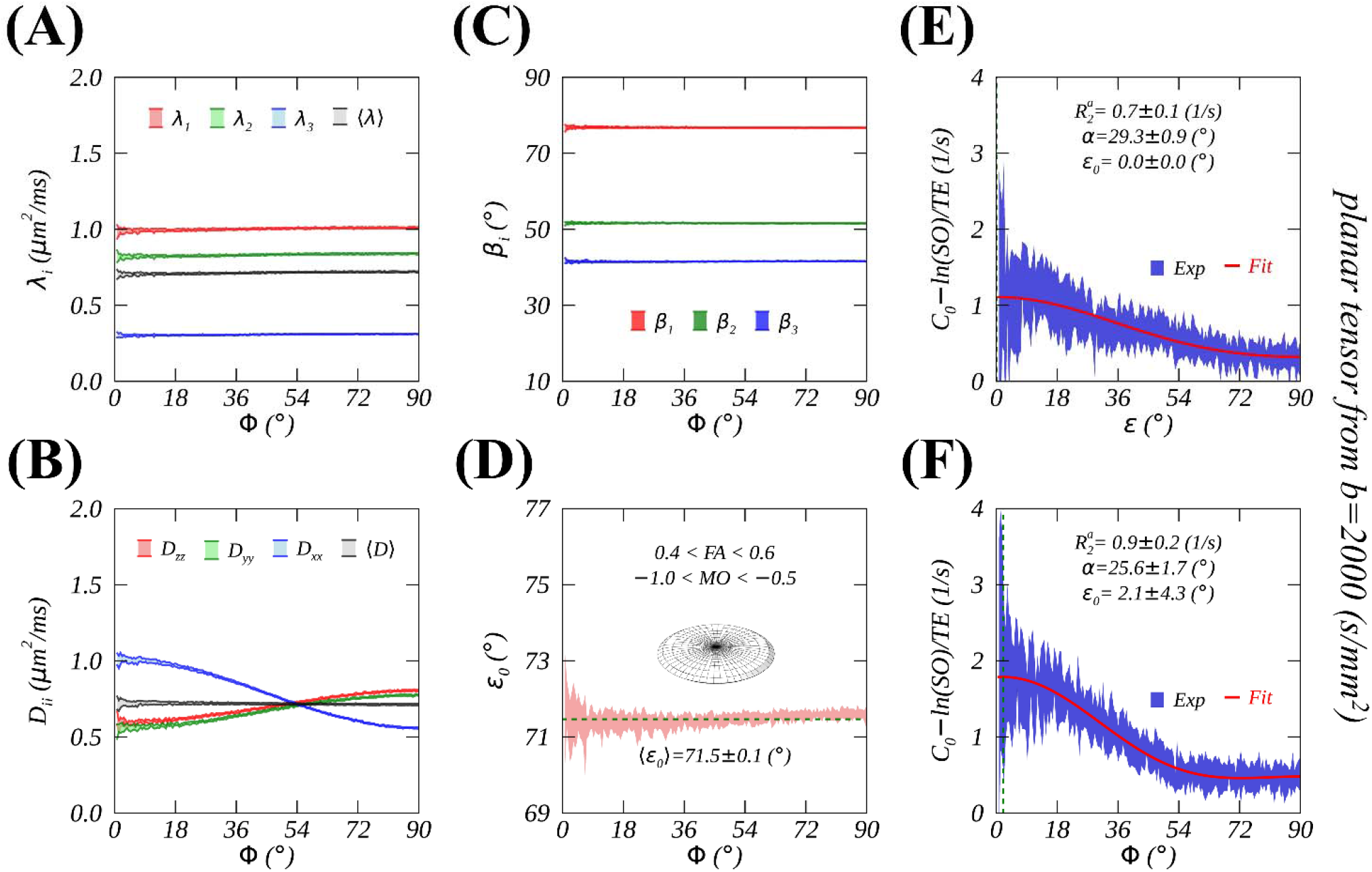
The same plots as Figure 5 except for the results derived from DTI data subsets (n=12) with *b*=2000 (s/mm^2^) containing planar tensor image voxels.

Compared to those from the linear tensor group, *R*_2_ orientation dependences from the planar tensor group manifest a 90° phase shift as predicted. Specifically, anisotropic *R*_2_ reached the minimum (e.g., Figure 6E) for a linear tensor group and the maximum (e.g., Figure 8E) for a planar tensor group when *ε*= 0°. Similarly, the fitted *α* from the planar tensor group was approximately complementary to that from the linear tensor group as tabulated in Table 2, i.e., 26° vs. 64°. Because of a smaller number of image voxels as demonstrated (i.e., ribbons in blue and black) in Figure 3F, relatively larger measurement uncertainties occurred in *R*_2_ orientation dependences, e.g., as shown in Figure 7F and Figure 8F.

## 5 DISCUSSION

This work has identified the origin of angle offsets manifested in previously reported *R*_2_ and 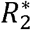 orientation dependences in WM. Diffusion data at two *b*-values from image voxels in both linear and planar tensor groups have validated theoretical predictions. The outcome from this work could have significant impact on the interpretation of past results and for future research.

### 5.1 An anisotropic diffusion direction

In this work, we suggest an anisotropic diffusion direction as a better representation for an axon fiber orientation in WM. This proposal was built on a well-established theoretical background^33^ and corroborated on both linear and planar tensors at two *b*-values. Our suggestion agrees well with what has been known from STI; more specifically, principal susceptibility direction always deviates from PDD based on DTI. It is not surprising when considering the facts that both anisotropic *R*_2_ relaxation and anisotropic susceptibility are induced by concentric distributions of rotationally restricted molecules (i.e., water and lipids, respectively) around an axon fiber in WM^46, 55^

The outcome from this work could have significant implications on interpreting clinical tractography in neuro applications or on evaluating quantitative MR results based on the standard orientation reference from DTI, e.g., microscopic susceptibility anisotropy imaging.^56^ Further, caution should be exercised when interpreting directional diffusivities (i.e., *D*_∥_ and *D*_⊥_) as a biomarker in axonal and myelin damage^57^ because they are not parallel to and perpendicular to an axon fiber as originally thought.

### 5.2 Diverse *R*_2_ orientation dependence patterns

The observed negative correlation between FA and *b*-values, as demonstrated in Figure 4E, had consequences on previously reported *R*_2_ orientation dependence patterns. These patterns guided by Φ had been shifted to a certain degree depending on *b*-values, e.g., Figure 6F vs. Figure 5F. Further, the measured anisotropic *R*_2_ amplitude (i.e., 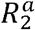) was relatively larger with a higher *b*-value. Generally, the FA threshold for classifying WM voxels is the same as done in this work regardless of varying *b*-values. Consequently, with a higher *b*-value, more image voxels with a lower FA would be disqualified for WM (see Figure 3E), leading to a relatively larger average FA for the selected voxels compared to those selected with a lower *b*-value. In short, with a fixed FA threshold, the selected WM voxels seem to be more anisotropic (i.e., an increased 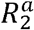) for higher *b*-values. Since no consensus on FA threshold and *b*-value was available for previous studies on anisotropic *R*_2_ and 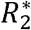 relaxation in WM, it is no surprise that some conflicting results could be found in literature.

### 5.3 A generalized MAE function

When an opening angle becomes zero, *f* (*α, ε*) in Eq. 7b will be shrunk into the familiar term (3cos^2^*ε* − 1)^2^ that is commonly associated with MAE in literature.^10, 16^ The measured anisotropic *R*_2_ or 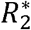 was larger with *ε* = 90° than that with *ε*= 0°, and the magic angle (i.e., *ε* = 54.74°) did not exist in WM, which seemingly justified the exclusion of MAE from being considered as a viable relaxation pathway.^9, 16, 20^ However, these conflicts between the theory and the measurement could be resolved when considering an axially symmetric distribution of bound water around an axon fiber as shown in this work.

The developed anisotropic *R*_2_ relaxation model is undoubtedly an overly simplified representation. According to an advanced Raman microscopy study on water orientation at the surface of phospholipid bilayers,^46^ an opening angle *α* of the funnel model should be equal to 90° for a single axon fiber while the fitted *α* was around 66° in adult WM. This clearly demonstrates that an additional source of bound water, with a uniform <H-H> distribution (i.e., *α* = 0), has contributed to the observed anisotropic *R*_2_ relaxation in WM. Based on previously reported data from several research groups (data not shown), the fitted appeared similar in adult WM but substantially decreased in neonate WM,^28, 43^ consistent with a declined FA.^28, 58^

### 5.4 An angle offset *ε*_0_ removal

As predicted, *R*_2_ orientation dependence became free from an angle offset *ε*_0_ when *ε* was used as an angle reference otherwise *ε*_0_ was nonzero as demonstrated in this work. When diffusion tensor was approximately axially symmetric, *ε*_0_ could be largely corrected by making an approximation on the least important eigenvalues for a linear tensor, i.e., *λ*_2_ ≈ *λ*_3_. This was not the case for a planar tensor, as shown in Figure 8D and 8F, where an approximation was made on the most important eigenvalues, i.e., *λ*_1_ ≈ *λ*_2_. If PDD had been utilized rather than TDD as used in this work, the angle offset *ε*_0_ would have been largely corrected as demonstrated lately.^45^ It is not an ultimate goal of this work to accurately correct *ε*_0_ using the predicated ⟨*ε*_0_⟩ that is inherently inaccurate. Rather, this planar tensor group provided a unique opportunity of the stringent test for the proposed theoretical framework, rendering not only the anisotropic *R*_2_ relaxation model but also the anisotropic diffusion model to be generalized for both single and crossing fibers in WM.

Water molecular rotational diffusion (related to *R*_2_) and translational diffusion (i.e., DTI) are intimately connected and thus have been collectively studied in recent years.^59^ To our knowledge, the present work is the first to bridge the most relevant degrees of freedom, i.e., an anisotropy,^60^ from the two different molecular motions for characterizing highly ordered microstructure in WM. The proposed theoretical framework makes it feasible to reliably assess to what extent myelin water imaging might depend on WM fiber orientation in the human brain,^25^ or to what degree anisotropic 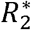 relaxation might hinge upon temperature and postmortem interval in a recent study.^24^

### 5.5 An unexpected orientation-dependent mean diffusivity

An interesting finding is that principal eigenvalue *λ*_l_ was somewhat orientationally variant, resulting in an angle-dependent mean diffusivity (i.e., black ribbon) as shown in Figure 5A. This unexpected variation decreased with a higher *b* value as seen in Figure 6A and almost vanished for a planar tensor (see Figures 7A and 8A) perhaps due to a limited SNR. Recently, a similar phenomenon was also reported by two research groups.^61, 62^ One group employed a lower *b*=700 (s/mm^2^) and found a slightly larger variation.^62^ Together with our observations with *b*=1000 and 2000 (s/mm^2^), it could be speculated that some interactions (or cross-terms) between imaging and diffusion sensitizing gradients might have played an important role in modulating mean diffusivity.^63^ Nonetheless, more research is warranted to further clarify this unresolved finding.

### 5.6 Limitations

There are several limitations to this work that deserve mention. First, a direct quantitative connection was missing between anisotropic rotational and translational diffusions except for the direction. Second, an isotropic *R*_2_ (i.e., 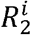), embedded in *C*_0_ in this work, was assumed invariant and thus cannot be determined. This shortcoming could be overcome using the conventional lengthy *R*_2_ mapping. 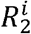 has been reportedly linked to iron deposit in WM, and an excessive iron deposit may play an important role in developing multiple sclerosis.^64^ Third, no effort was made to determine to what extent the susceptibility effect had contributed to the reported orientation-dependent *R*_2_. Nonetheless, we speculate that this effect might have played an insignificant role at 3T because *R*_2_ was known to be relatively unchanged either at low (e.g., 1.5T) *B*_0_ fields in neonate WM^65^ or at ultralow (e.g., 50mT) *B*_0_ fields in adult WM.^66, 67^ A recent study on 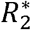 orientational anisotropy at 3T and 7T provided direct evidence to support our perspective in which an increase in 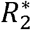 anisotropy from 3T to 7T was much less than expected if the susceptibility effect was considered alone.^68^ Last, the reported *R*_2_ orientational anisotropy is non-local in nature and thus absent is the voxel-wise ultrastructural information. An intrinsic orientation-independent metric for each image voxel could be deduced from *R*_l*ρ*_ dispersion imaging^30^ as recently demonstrated on articular cartilage.^32, 69^

## 6 CONCLUSIONS

In summary, we have identified the origin of *R*_2_ and 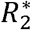 orientation dependence angle offsets and demonstrated an anisotropic diffusivity direction as a better representation for orientations of both single and crossing axon fibers. Further, the developed generalized magic angle effect function rendered full quantification of *R*_2_ and 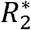 orientational anisotropies, thus shedding new light on microstructural alterations in the human brain WM.

## ACKNOWLEDGEMENTS

The author would like to thank Prof. Thomas L. Chenevert at University of Michigan for many useful discussions on diffusion imaging and Dr. Peter J. Basser at NIH for his support and encouragement.

## DATA AVAILABILITY STATEMENT

Data used in this work are available on request from the author.

